# Single particle trajectory statistic to reconstruct chromatin organization and dynamics

**DOI:** 10.1101/559369

**Authors:** O. Shukron, A. Seeber, A. Amitai, D. Holcman

**Author notes:** equally contributing.

## Abstract

Chromatin organization remains complex and far from understood. We discuss here recent statistical methods to extract biophysical parameters from in vivo single particle trajectories of loci to reconstruct chromatin reorganization in response to cellular stress such as DNA damages. We look at the methods to analyze both single loci as well as multiple loci tracked simultaneously and explain how to quantify and describe chromatin motion using a combination of extractable parameters. These parameters can be converted into information about chromatin dynamics and function. Furthermore, we discuss how the time scale of recurrent motion of a locus can be extracted and converted into local chromatin dynamics. We also discuss the effect of various sampling rates on the estimated parameters. Finally, we discuss polymer methods based on cross-linkers that account for minimal loop constraints hidden in tracked loci, that reveal chromatin organization at the 250*nm* spatial scale. We list and refer to some algorithm packages that are now publicly available. To conclude, chromatin organization and dynamics at hundreds of nanometers can be reconstructed from locus trajectories and predicted based on polymer models.

## 1 Introduction

Fundamental physical properties of DNA such as the internal elasticity, the bending properties as well as the rotational energy have been estimated ex-vivo [66, 67]. But what about the properties of chromatin in vivo? Not only was it simply difficult to visualize specific chromatin loci but it remains unclear to date, what exact physical quantities should be measured. Shall we simply use the same classical physical observables measured ex vivo or shall we focus on other parameters to be extracted so that we can hope to reconstruct from the space a locus explores the chromatin dynamics? Observables are physical quantities that can be measured but they depend on the perception of what we are studying and a priori model of the phenomena. This choice made by the observer is thus open to bias. In the case of chromatin, we still do not have a complete and definitive physical model that would represent its dynamics and possible configurations. Recent works by us and others have brought chromatin models forward to the point that they can be predictive [28, 7] for its organization and dynamics. Here we present studies combining single particle trajectories (SPTs) of one and multiple chromatin loci with polymer modeling that aim to clarify chromatin structure and dynamics and how it changes in response to DNA damages.

Modeling chromatin with polymers started with the primitive Rouse model (beads connected by spring), on which repulsion forces have later been added (Lennard-Jones forces) followed by bending elasticity, reviewed in [7] (Box 1). Recently, to account for long-range forces between various parts of the chromatin, connecting monomers located far away have been introduced in simulations [10, 29, 9] and theory [5, 60]. Although these models are used to simulate chromatin in some defined conditions, how they relate to the analogous biological condition is unclear. To connect polymer models and how chromatin actually behaves inside the nucleus one must be able to extract physical parameters from real experimental chromatin data under various conditions. To this end, the adoption of green fluorescent protein (GFP) and other fluorophores to visualize single chromosomal loci in vivo [12] enabled many genomic loci, under multiple conditions to be studied [57]. This led to the advancement of several biological questions regarding the cellular and genetic components that regulate chromatin movement, such as cell cycle stage and cohesin loading [30, 17] or the role of nucleosome remodeling enzymes such as the INO80 complex [50], reviewed in [16]. The function of chromatin movement is not well understood and is difficult to experimentally prove as many of the factors that regulate movement also have other roles in the cell. One role in which chromatin movement may contribute, is the efficiency of repair of DNA double-stand breaks (DSBs) by a process called homologous recombination HR, reviewed in [28]. Here a broken DNA strand must physically scan the nucleus to find its homologous partner if the replicated sister is not present or is otherwise damaged. While definitive proof that chromatin motion facilitates repair by HR remains elusive, there are a number of correlations. Ablation of proteins that result in reduced movement, decreases repair efficient by HR [28], while mutations that increase movement improve HR efficiency [50, 58]. However, while we know that chromatin movement is regulated in some conditions, we understand little about how chromatin moves. Limiting our understanding is how single particle trajectories are generated as well as how they are analyzed.

This paper is organized as follows: The first section describes physical models that have been used to analyze chromatin trajectories. We introduce four key physical parameters that can be extracted from chromatin loci trajectories and can be used in combination to characterize motion. The second section presents how to build a correlation analysis for two loci recorded simultaneously. In the third section we discuss crowding constraints on chromatin, polymer simulations and what can be predicted about chromatin motion. Finally, we summarize existing publicly available algorithm packages. Although many examples described here concern yeast, the statistical methods and data analysis can be applied to many other organisms, from insects to mammalian cells [69, 47, 13].

## 2 Chromatin locus dynamics revealed by SPTs

### 2.1 Which model to use for SPTs: free, confined or anomalous diffusion?

SPTs consist of an ensemble of successive points acquired at a sampling rate Δ*t*. What can be extracted from these trajectories? To answer this question a physical model, which is a framework that provides plausible physical mechanisms and accurate predictions, should be selected. However, this selection process remains challenging. For example, in the case of the nucleus, finding the best physical description of chromatin will provide an optimal framework to define, elucidate and predict its motion in various conditions. We now present briefly some elementary scenarios to infer the underlying physical processes based on trajectories.

The most well-known example of molecular dynamics not driven by any active mechanism is classical diffusion (see Glossary) described by Brown in 1827 and quantified statistically by Einstein in 1905. A particle driven by free diffusion at scale Δ*t* is characterized by random jumps that follow a Gaussian distribution of variance 2*D*Δ*t*, where *D* is the diffusion coefficient. In that case, the physical quantity to estimate is simply the diffusion constant *D*. However, the locus motion can be restricted due to obstacles or to some resulting tethering force, which are not accounted in classical Brownian motion. In that case, how can we determine the extent of the restricted region and the nature of the restriction (confinement vs tethering force)? We will disucss below some associated parameters to be measured that allow for us to distinguish these models.

Another possibility to describe the motion of a locus is to use a deviation from classical Brownian motion, which is call anomalous diffusion, where the forces underlying the locus dynamics are correlated in time [49]. In that case, such correlation should be determined. Another possibility is that SPTs can result from a combination of deterministic (generated by a force) and Brownian motion. The motion could also result from switching between a deterministic forces and diffusion. In that case, the nature of the force should be identified as well as the switching rate, often approximated as Poissonnian (characterized by a single exponential parameter). Finally, we recall that the trajectory of a tagged locus can only provide a local and partial information about the long-range chromatin interactions. Thus, polymer models are used to integrate this local information into a general framework.

To summarize, we need to select among various possible physical models, making an informed decision based on trajectory data. Thus, we are left with the following question: how to infer from the motion of a locus, the optimal polymer model that describes chromatin? When successful, the underlying model can be used to extract specific parameters that characterize chromatin under different conditions: transcription, DSBs, DNA viral translocation, various drugs, different cellular phases, etc. such approach is now currently used to study chromatin re-organization.

### 2.2 From trajectory analysis to model selection

In this section, we describe four parameters computed from SPTs that are used to select one of the physical model for chromatin dynamics mentioned in the previous subsection. If the locus is assumed to be diffusing in a free space, then free diffusion is characterized by a single parameter: the diffusion coefficient *D*_*c*_, which should be estimated from data (see below). For confined and restricted motion, more parameters are needed to determine the restriction. The restriction could be due to the external environment such as crowding but not only: spatial restriction of chromatin in special configurations, such as the Rabl configuration of budding yeast chromosomes, where telomeres and centromeres are tethered to the nuclear periphery [68], will limit chromatin motion. Therefore, the diffusion coefficient of a chromatin locus is clearly insufficient to describe confinement.

#### 2.2.1 Diffusion coefficient

The first step is to collect the data and then the diffusion coefficient can be estimated along trajectories (see Box 2). Computing and comparing the diffusion coefficient estimated at different locations clarifies the spatial heterogeneity within the nucleus. Interestingly, the same loci in different cells can have different diffusion coefficient, demonstrating the variability across cells, the reason for which remains unclear.

While extracting the diffusion coefficient *D*_*c*_ of a single chromatin locus is limited due to many factors affecting the statistical evaluation of this variable, it is possible to extract the diffusion coefficient from many different trajectories. Indeed, instead of computing the diffusion coefficient along trajectories, it is possible to combine a massive number (10^4^ – 10^5^) of super-resolution trajectories to reconstruct a diffusion map inside cellular environments, membranes and the nucleus for a variety of fluorescently tagged proteins such as histones, receptors, cytosolic proteins [48, 22, 21, 35, 23, 40, 46, 36, 32, 38, 33].

#### 2.2.2 Anomalous exponent

The anomalous diffusion is a deviation from classical diffusion, characterized empirically by an anomalous exponent *α* that can be extracted from the slope of the Mean Square Displacement (MSD) 〈|*X*(*t* + Δ*t*) − *X*(*t*)|^2^〉 that behaves like *At*^*α*^ for Δ*t* small over the time (Box 2). When *α* = 1 it reflects a Brownian motion, *α* > 1 is called super-diffusion, which represents a stochastic dynamics containing a deterministic directed motion. Finally, *α* < 1 is a sub-diffusive motion, which could result from short-range forces (see below) or viscosity memory [42, 5, 43, 7].

Note that directed motion is often hidden in the MSD analysis. In that case, extracting the anomalous exponent is not enough to account for the entire dynamics. In parallel, a drift analysis has to be performed to extract the distribution, organization of vector fields, [35, 39, 34] and directional changes [11, 52].

The exponent *α* computed for a single locus remains difficult to interpret and in particular to connect with local chromatin organization. *α* reveals much of the nature of a locus that belongs to a polymer, but clearly remain insufficient, alone, to recover all the properties of chromatin. In general, the exponent *α* for chromatin loci varies in the range 0.3-0.5, when no external forces are applied to the polymer model [7]. But in the presence of deterministic forces, generated for example by the nucleus surface oscillation, actin or microtubule network, the range of *α* can increase to (0.5 − 2).

#### 2.2.3 Confined diffusion

How much space a locus explores in a given time is another important parameter to extract from a trajectory. To quantify spatial restriction due to crowding or chromatin anchorage, the length of constraint *L*_*c*_ can be used [7]: It is defined as the standard-deviation of the locus position with respect to its mean averaged over time (see Box 2). This parameter provides an estimation for the apparent radius of the volume explored by a trajectory and importantly does not require a plateauing of the MSD. A small *L*_*c*_ compared to the radius of the nucleus is thus the signature of a high confinement, while a large *L*_*c*_ means that the locus motion is not restricted by any nuclear bodies or chromatin. In summary, the length of constraint *L*_*c*_ provides a measure of confinement but it does not reveal the underlying mechanism.

#### 2.2.4 Tethering force to restrict diffusion

What restricts the motion of chromatin loci? A trajectory could be confined either by obstacles or by a field of force. How can we differentiate between these two possibilities? This question can be addressed by using a polymer model, where a tethering force is generated by a potential well 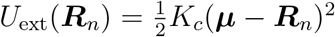 centered at a point ***μ*** and applied at a monomer *R*_*n*_. The spring constant *K*_*c*_ has to be estimated from data (Fig. 1A and Box 2). The field of force *U*_ext_ should influence the motion of any locus and thus could be extracted from trajectories. This tethering not only affect the motion of the whole polymer (chromosome) but can arise from interactions of the locus with other chromosomes or nuclear substructures such as the nuclear envelope. While it is not possible to directly measure this force, it can be inferred from SPTs [3]. Indeed, a procedure has been developed for freely moving particle [38] requiring many redundant trajectories exploring the same micro-environment. However, for a single-long trajectory that does not come back too many times on itself, there may be not necessarily enough data points. To overcome this difficulty, a recent method [3] considers consecutive displacements *R*_*c*_(*t*+Δ*t*)−*R*_*c*_(*t*) as independent when a trajectory is recurrent returning back on itself many times and in that case, it is possible to estimate the spring constant *K*_*c*_, as described in Box 2 and [4]. To conclude, when a trajectory folds back on itself many times, much like a spring, it is possible to extract a field of force [73].

**Figure 1:**
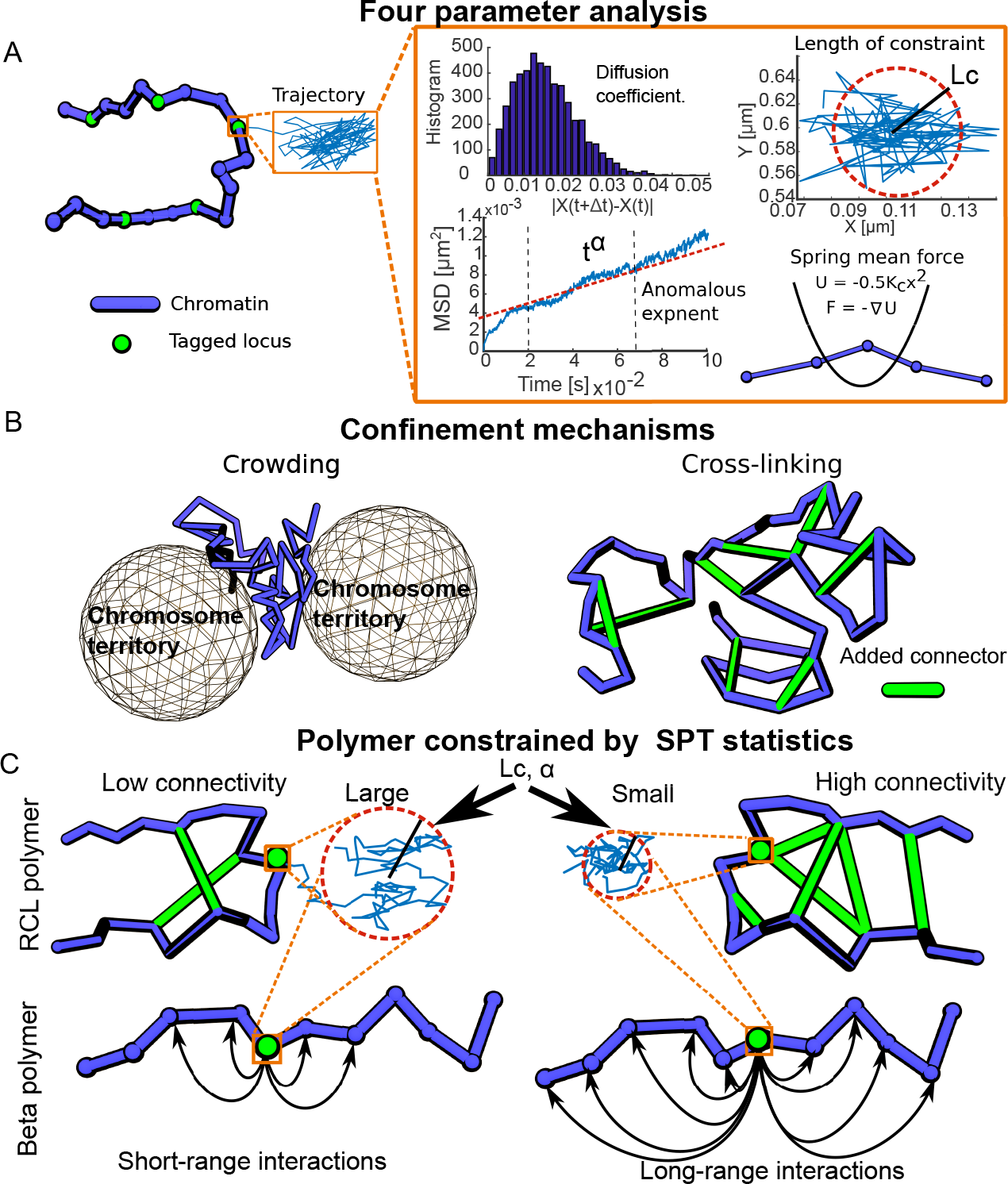
From SPT of chromatin locus to polymer models. **A.** Extraction of four parameters from a trajectory of a single locus (green). The four parameters extracted are: the diffusion coefficient *D*_*c*_, the length of constraint *Lc*, the anomalous exponent *α*, and the spring constant *Kc* of a parabolic potential well (red box). **B.** Two possible confinement mechanisms: crowding can be generated from the exclusion of a polymer from chromosome territories, thus leading to the restriction of a locus (left), or confinement can be induced by added cross-linkers (such as CTCF, cohesin, condensin) to the chromatin fiber. **C.** Construction of polymer models constrained by SPT statistics: values of parameters such as *Lc* or *α* can be used to constrain polymer such as the RCL polymer (top) or the beta polymer (bottom). For example, *Lc* and *α* small can be recovered by adding the right number of connectors (green) or long-range interactions (arrows) to RCL and beta polymer model, respectively.

The field of force discussed above reflects the internal chromatin interaction as well as the external forces due to long-range interactions between genomically distant loci that may be are brought together spatially by CTCF or cohesin. While budding yeast do not have CTCF they do have cohesin, which holds sister chromatids together in S phase. Artificial removal of cohesin in S phase cells increases chromatin motion. Interestingly, removal of cohesin in G1 phase cells, where the sister chromosome is absent, also marginally increases chromatin motion. This suggests a role for cohesin as a possible connector in budding yeast [70, 17].

For motion confined due to crowding only, applying the above procedure would not lead to any forces (which means *K*_*c*_ = 0), because there is no field of force, except for the points in very close vicinity of the large obstacles [35] that restrict motion (Fig. 1B). When obstacles appear clearly, such as the rDNA containing nucleolus or the boundary of the nucleus, these estimations work well, but when obstacles are completely intermingled, these estimations are less robust.

To conclude, the four parameters described above (*D*_*c*_, *α*, *L*_*c*_, *K*_*c*_) can be estimated from trajectories and they provide complimentary information about the statistical properties of chromatin loci [28, 7]. In addition, their estimation does not assume or require any underlying physical structure such as a polymer model, which will come in a second stage for the interpretation and the reconstruction of the underlying chromatin organization at a given spatial and temporal scale (see Box 1).

### 2.3 Relating polymer models and the anomalous exponent

How to chose a polymer model from the statistics of SPTs? We recall that for Rouse polymer, consisting of monomers connected by spring the anomalous exponent for each monomer (except the extremities), the anomalous exponent is *α* = 0.5, already illustrating a clear deviant behavior compared to Brownian motion (for which *α* = 1). This difference in *α* shows how polymer dynamics influences the motion of a single locus. Recently, a new class of polymer model was introduced (the *β*-polymer model) based on adding mid and long-range forces between all monomers to be calculated specifically [5] for anomalous exponents in the range of *α* ∈ [0 – 0.5]. For example, let us choose *α* = 0.345, what is the exact distribution of forces (spring constant) between monomers that leads to this value? This distribution was computed in [5] and could be seen as a realization of chromatin with cross-linker molecules. This model works for *α* greater than 0.5 and has been used to interpret the motion of DSBs in budding yeast [28, 7]. Here, after removal of a deterministic force that masked changes in *α*, an increase was observed at the site of a DSB. But what does an increase in *α* mean? While it could indicate that locus starts to move with more directed motion, it tells us little about the structure of the locus we observe or its interactions. However, using the *β*-polymer model and numerical simulations, an expansion of chromatin at the site of a DSB was predicted when *α* increased. This prediction was confirmed by super-resolution microscopy [7]. In this case the polymer model was driving a testable hypothesis and led to the design experiments that confirmed the theoretical prediction.

Reconstructing chromatin when the motion of loci is characterized by *α* > 0.5 is more complicated, as different type of forces are needed: combining a *β*-polymer model with a deterministic drift, which can even be oscillatory [7] increases the anomalous exponents. In addition to the *β*–polymer model, there are other ways to study long-range interactions in chromatin. One way is to use a model were the monomers of a polymer are connected randomly with transient (Binder model or RLM theory [10, 9]) or fixed (RCL theory [60, 63]) connectors (Fig. 1C). These scenarios lead to an anomalous exponent for each locus < 0.5, although the exact relation between a realization characterized by the distribution and the number of connectors, and the anomalous exponent remains unclear.

Adding connectors to a polymer leads to confined motion of its monomers (Fig. 1B). From SPTs of a single locus, it is possible to extract a resulting force only. It thus remains an open question how to reconstruct from such statistics the exact short, mid and long-range (materialized by binders such as CTCF) forces that need to be added on a polymer model. However, extracting forces that restrict motion is very accurate from tens of thousands of super-resolution SPTs, leading to the reconstruction of a drift map [39]. In the case of long SPTs, it is still possible to estimate forces from the recurrent motion [4]. The reconstruction assumes that confinement results from a parabolic potential, as discussed in subsection 2.2.4 and Box 2. This is a classical model used in chemical reaction and characterised by a finite parabolic well, truncated at a specific height. Recovering these characteristics is possible from many trajectories [33].

For the chromatin, the story does not stop here. Once an overall force is computed, due to the nature of the polymer, a second step is needed that consists in a deconvolution procedure [4] which is required to differentiate the internal (due to chromatin internal properties) from external forces. This analysis is performed under several assumptions and when the chromatin is approximated as a Rouse polymer model, it is possible to recover the spring constant of the effective external force from the total force, when we know the genomic distance between the tagged locus and the one where the force is applied to (Box 2). The case of multiple forces was resolved in the Supplementary Info of [4]. For more elaborate models, numerical simulations are used to recover external forces. To conclude, the origin of forces acting on chromatin is still poorly understood, but polymer models suggest that connectors restrict chromatin loci motion, leading to apparent forces. The amplitude of these forces has been used to quantify chromatin dynamics before and after induction of DSBs [7, 28].

How to interpret changes in the anomalous exponent *α*? Such interpretation depends also on the polymer model: changing the number of connectors or the forces between monomers in the *β* or RCL polymer respectively modifies *α*. In these models, a decrease in the anomalous exponent is always obtained by local chromatin compaction, while an increase is associated to decompaction [7]. A decrease in the anomalous exponent *α* is associated with an increase in the mid and/or long-range forces between monomers, leading to chromatin compaction. Conversely, an increase in *α* is associated with force releases, leading to chromatin decompaction. These theoretical results led to the prediction that after the induction of a DSB, chromatin decondensed locally, and this was interpreted as release of long-range forces between chromatin loci. A possible scenario could be that removing or adding random connectors (Fig. 1C), such as cohesin and/or CTCF modulate chromatin organization. To conclude, short and long-range local chromatin organization influences SPTs of a single locus and can be estimated from *α*.

Another possible scenario for confinement comes from random cross-linkers [60] between chromatin loci (Fig. 1B-C). This hypothesis can be evaluated by looking for simultaneous changes of the four parameters presented above: for example, a reduction of the length of constraint *L*_*c*_ in parallel with a reduction of the anomalous exponent *α* and an increase in the local tethering force spring *k*_*c*_ is associated with an increase in the number of connectors and chromatin condensation. However, no changes are expected to be seen in the diffusion coefficient [7], unless there is a local nucleosome removal at the measured site. Conversely, a decrease of the diffusion coefficient is directly connected to an increase of crowding [31] obtained by adding sufficiently large obstacles compared to the size of the observed loci [63].

To conclude, changing the number of connectors between chromatin loci has several consequences: First, only a few connectors are required to see a change on polymer dynamics. In general, less than 1-5% are sufficient to restrict chromatin motion, but as discussed below, many more are needed to guarantee a radius of gyration of a few hundred nanometers. Second, connectors can be positioned at random and not at a specific location, leading to a spectrum of anomalous exponents computed at each loci. The number of connectors can be estimated using the encounter probability of the HiC data [63]. This number seems to play a key role in defining topologically associating domains (TADs) [63], gene regulation [63] and organization across cell differentiation [62].

### 2.4 Influence of the sampling time step Δ*t* on the four measured parameters

How does the sampling time Δ*t* influence the biophysical parameters extracted from loci SPTs? First Δ*t* enters directly in the estimation of the four parameters mentioned above (Box 2). Second, from polymer model theory, when Δ*t* is small or large compared to the relaxation time scale [18, 5, 60] of the chromatin, we expect to extract difference in parameters such as diffusion coefficient or the anomalous exponent *α* [6]. In general there is no simple relation between the time step Δ*t* and the underlying physical property of chromatin and its interaction with the environment (Fig. 2A). For example, can you interpret that decreasing Δ*t* is associated with an increase of the effective diffusion coefficient? For a random walk (Brownian motion at scale Δ*t*), changing Δ*t* has no effect on estimating the diffusion coefficient *D*_*c*_. However, for chromatin, decreasing the sampling rate Δ*t* increases the effective diffusion coefficient [7]. One possible interpretation is that by increasing the time step, more of the chromatin contributes to the motion of the locus, leading on average to a decrease in the effective diffusion coefficient (Fig. 2B table). Conversely, by decreasing the time step Δ*t*, the diffusion coefficient *D*_*c*_ increases because the part of the polymer influencing the motion has diminished, leading to a reduction in the resulting estimated force and thus less constraints are influencing the monomer motion. The choice of Δ*t* is thus critical: it should not be too small (that would only capture the motion of a single monomer) or not too large (when the entire polymer can move by diffusion in a confined environment), but be chosen in an intermediate time regime, as predicted from the anomalous behavior of a monomer [5, 7].

**Figure 2:**
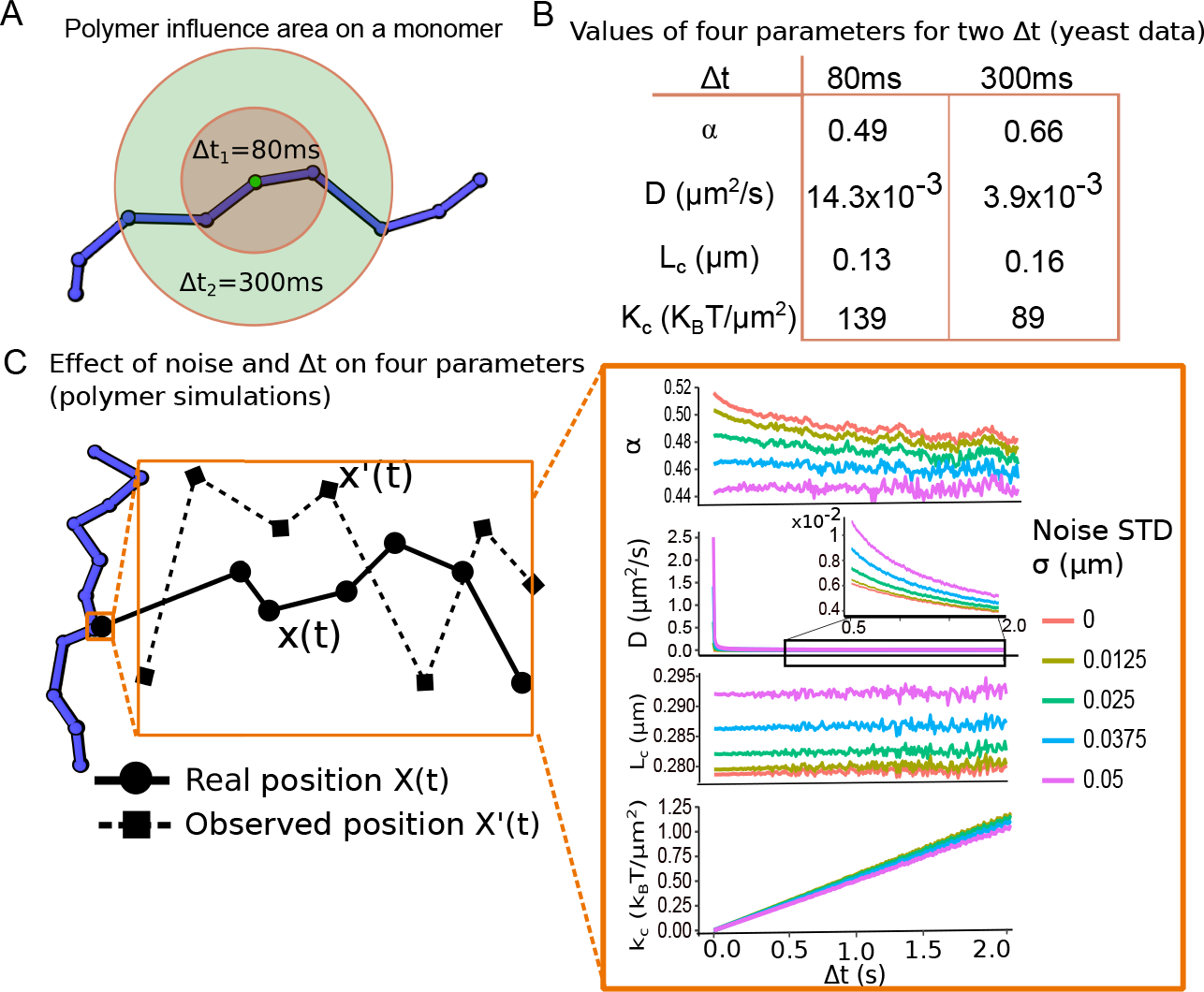
Influence of the sampling rate Δ*t* and localization noise on SPT statistics. **A.** The region of influence on a locus (green) depends on the sampling rate Δ*t*: as Δ*t* increases the fraction of the polymer influencing the motion of a single locus increases, at 80ms (brown) the region is smaller than the one obtained at 300ms (green). **B.** Example of changes in the four parameters with respect to the sampling rate Δ*t*, obtained from yeast SPTs [7]. **C.**Influence of the localization noise on SPTs. The localization noise generates a Gaussin error on the position of loci at each time step Δ*t*. Note, this noise can affect the four parameters: *α*, *D*_*c*_, *Lc*, *Kc*.

Another effect of Δ*t* concerns the interpretation for the tracking localization noise (Fig. 2C), where a Gaussian error of amplitude *σ* is added to the physical position *Z*_*phys*_, so that the measured position is *Z*_*measured*_ = *Z*_*phys*_ + *σξ*. Such localization error affects the value of the effective diffusion coefficient compared to the physical diffusion coefficient *D*_*phys*_, leading to shift 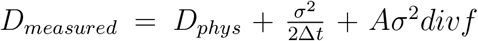, where A is a constant and *f* is the force applied to the tagged loci [37]. The noise localization error influences differently the values of the four biophysical parameters, in particular, the anomalous exponent *α* should not vary when Δ*t* changes because 〈|*Z*_*measured*_(*t* + Δ*t*) − *Z*_*measured*_(*t*)|^2^〉 ≈ *C*(Δ*t*)^*α*^ + *C*_2_, where *C*_2_ is a constant. However, experimental results shows that it is an increasing function of Δ*t*, suggesting that other physical mechanisms should be taken into account when increasing the time step Δ*t*.

Interestingly, the diffusion coefficient can also depend on Δ*t*. Indeed, for small Δ*t*, obstacles will restrict motion and the estimated diffusion coefficient (Box 2) will only capture the space between obstacles. However, increasing Δ*t*, another diffusion regime (effective diffusion) can be captured, which consists in large random jumps between obstacles. For example, observing a change by a factor two in the effective diffusion coefficient is associated with an increase of the density of obstacles from 50 to 70% [31].

To conclude, it may appear contradictory to extract at the same time a diffusion coefficient, assuming the underlying process is a classical diffusion and the anomalous exponent, referring to an anomalous description. These two descriptions could co-exist: trajectories could be seen as generated by an apparent diffusion process at a fixed sampling time scale Δ*t*, but also by an anomalous process, coarse-grained at a scale Δ*t*. The time step is certainly critical for the parameter estimations. Future investigations are needed to clarify what exactly can be revealed from the chromatin at small and large Δ*t*.

## 3 A first passage time analysis of two simultaneous tracked loci reveals chromatin dynamics at the 250nm scale

Are there new features or biophysical information contained in trajectories of two loci tracked simultaneously that are not contained in loci tracked individually? Using the four biophysical parameters mentioned above, each locus could be analyzed separately. However, we shall discuss here how the statistics of the distance changes between the two loci reveals chromatin organization and dynamics at a scale of hundreds of nanometers [45, 61] for the analysis of the data of [15, 28]). We now recall a method introduced in [61], which focuses on the recursion dynamics of the two loci and the distribution of times *τ*_*D*_ and *τ*_*E*_ that they are spending in closed proximity and far away before they meet again, respectively. Intuitively, the distribution of these two times reveals the local organization of the chromatin: the time *τ*_*E*_ to meet again is associated with the local organization in a scale of the order of the genomic distance between the two loci, while the time *τ*_*D*_ accounts for the local three dimensional organization around the two loci.

The method of analysis has several steps: The first one is to consider the two trajectories *X*(*t*), *Y* (*t*) (Fig. 3A) and an arbitrary threshold length *T* = *ɛ*, that defines an encounter distance, which we can vary between tens to hundreds of nanometers. The second step is to compute the distance *d*(*t*) = |*X*(*t*) − *Y* (*t*)|. Then the segmentation consists in collecting the time interval before the two locus encounter for the first time, defined by *d*(*t*) > *ɛ*, leading to a time series 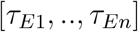 and a second time series for the distribution of times before the two loci dissociate 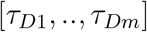 (Fig. 3B). In the third step, the two time distributions are well approximated by an exponential (Fig. 3C). This property is valid for several genomic distances between the two loci [61]. The exponential decay of the encounter time distribution is well predicted in the context of rare events of polymer looping theory [53, 2, 6]. If the loci dissociate within a physical mechanism against a field force (that tends to prevent the two loci from separating), then the dissociation time distribution can also be approximated by a single exponential distribution.

**Figure 3:**
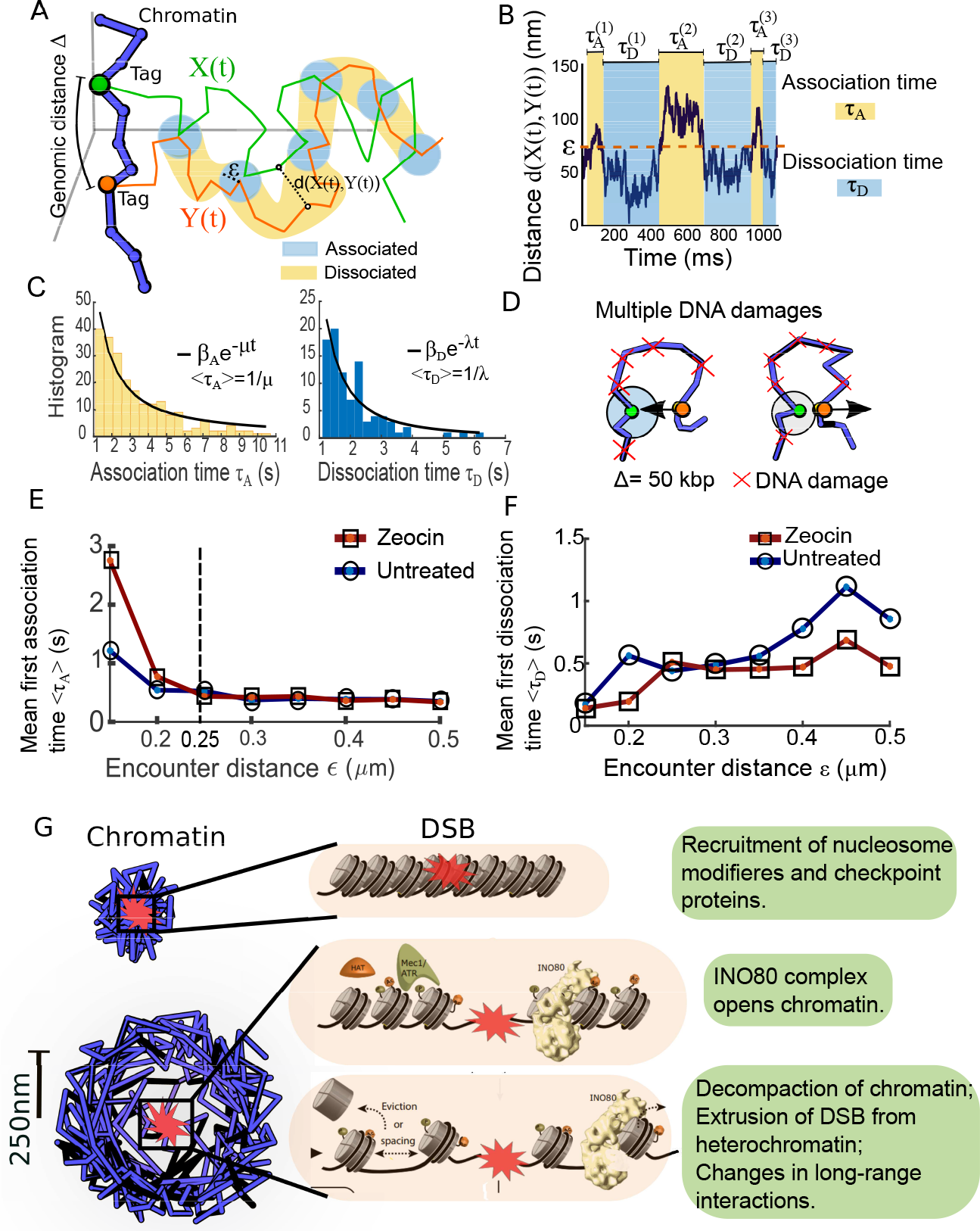
Multiple loci SPT statistics. **A.** Two loci (green, orange), separated by a genomic distance Δ, generate two trajectories *X*(*t*), *Y* (*t*), showing regions where they are located inside balls of radius *ϵ* (blue) separated by regions (yellow) where the trajectories are far apart. **B.** Distance *D*(*X*)*t*, *Y* (*t*)) versus time. When *D*(*X*(*t*), *Y* (*t*)) is lower than a threshold *ϵ*, trajectories are classified as associated (blue, Panel A) and characterized by the time (dissociation time *τ*_*D*_) it takes for the two trajectories to separate. Conversely, when *D*(*X*(*t*), *Y* (*t*)) > *ϵ*, trajectories are dissociated (yellow, Panel A), and characterized by the time (association time *τ*_*A*_) it takes for the two trajectories to enter for the first time into a region of radius ≤ *ϵ*. **C.** Distribution of association and dissociation time, characterized by a single exponential. **D.** Schematic representation of two loci association (left) and dissociation (right) dynamics with multiple DNA damages (red crosses). **E.** Mean first association time 〈*τ*_*A*_〉 versus the encounter distance *ϵ* for zeocin (red, squares) and untreated (blue, circles) chromatin. **F.**Mean first dissociation time 〈*τ*_*D*_〉 versus the encounter distance *ϵ* for zeocin (red, squares) and untreated (blue, circles) chromatin. **G.** consequences of DNA damages of local chromatin reorganization. Material is extruded following DSB.

This passage time analysis was applied initially to two loci SPTs [15, 28], where the distance between the two tagged loci varied in the range [23.5, 100.8] kbps and were tracked at time intervals of 300ms over 60s. The mean encounter time 〈*τ*_*E*_〉 decreases in the range *ɛ* ∈ [0, 0.25]*μm*, but was independent of the encounter distance for *ɛ* > 0.25*μm*. In comparison, the mean dissociation time 〈*τ*_*D*_〉 increases with *ɛ* in the range [0, 0.5]*μm*. Later, the analysis was applied to two loci located at a distance Δ = 50*kbp* apart tracked over 60 – 120*s* at 300ms rate, before and after the use of the drug Zeocin 500*μg/ml*, which induces DNA damages at random positions of the chromatin (Fig. 3D, top, red crosses). The analysis revealed that the mean duration 〈*τ*_*E*_〉 decreases with respect to the distance threshold *ɛ*, both in the untreated and Zeocin treated cases (Fig. 3D): a plateau is reached in 500ms for *ɛ* = 0.25*μm*, showing that the encounter time does not depend on *ɛ* above 0.25*μm*. In addition, the mean encounter time 〈*τ*_*A*_〉 (Fig. 3E) is slower following induction of a DSB, while the mean dissociation time 〈*τ*_*D*_〉 is faster (Fig. 3F): this result suggested that chromatin is less crowded at the spatial scale of the order of the distance between the two loci, while it is more crowded far away. This may be due to a local depletion of material locally after DSB, which is relocated further away from the break site.

To conclude, distributed DSBs impair recurrent chromatin motions only at a scale below 0.25*μm*, suggesting that this spatial scale characterizes the local chromatin organization in which undamaged loci can freely move, but become restricted above it (Fig. 3G). We expect that recording multiple loci at the same time could reveal the local chromatin organization and could be used to determine the minimum number of cross-linkers compatible with the statistics of the recurrent loci behavior.

## 4 Local chromatin environment reconstructed from simulations of polymer model

Understanding how chromatin moves in a confined environment remains challenging, but particularly useful to analyze the local environment located near a DSB. What can we learn about very confined chromatin environment from numerical simulations? The first approach to answer this question is to identify a polymer model and to constrain it by the four biophysical parameters mentioned in section 2 and extracted from SPTs.

### 4.1 Cross-linkers constrain chromatin motion

We discuss here evidences of short and long-range interactions of cross-linkers that constraint chromatin motion. Indeed, there are around ~100,000 CTCF binding sites identified in the genome[44, 56], but not all binding sites are occupied at any moment of time. Genomic loops mediated by structural proteins vary in the range of 15-123kbp in HeLa cells, with a mean of 86kbp [41]. A significantly larger cohesin loop of ~ 370 kbp was measured in [26]. Bound cohesin is not distributed uniformly along the chromosome, having higher concentration near centromeres and overall average spacing of 10kb apart [65]. In addition, cohesin binding sites correlate with position of DNA replication origins [25].

Although CTCFs, cohesin and condensin molecules are bound to chromatin, it is still unclear how exactly they are participating to loop formation. But CTCF residence time is ~ 1min in mouse embryonic stem cells [27]. Approximately ~80% of CTCF molecules bind transiently and non-specifically to the chromatin with a residence time of ~0.2-0.6s [1], whereas the residence time for a residual fraction of CTCF is in a wide range between ~4s to > 15min. Residence times for cohesin are longer than those of CTCF, and are cell cycle dependent. In G1, about 30% of cohesin is bound to chromatin with residence of ~6 hours, whereas in G2 about 45% of cohesin is bound with a residence time of ~24 min, as measured in [24]. Similar times of ~20min were also measured in [27].

In that context, two type of models have been used: those with transient [10, 9] and the others with permanent but randomly positioned connectors [63, 60]. The local chromatin environment including constraints can be simulated using polymer models such as the *β*-polymer or Randomly Cross Linked (RCL) model [63, 60]. The later model describes chromatin containing cross-linkers, positioned at random places. We recall that the RCL polymer is composed of a linear Rouse backbone of *N* monomers, connected sequentially by harmonic springs. In each realization of the polymer, *N*_*c*_ spring connectors are added (Fig. 4A) between randomly chosen non-nearest monomers pairs. This parsimonious addition of connectors accounts for binding molecules such as CTCF and Cohesin or Condensin, and their random position can generate the heterogeneity in chromatin architecture observed across cell population. The key parameter that reflects constraint of the chromatin encounter probability is the number of connectors in a defined sub-chromatin region, but not necessarily their position, as long as they are uniformly distributed. Transient and static connectors lead to similar statistical properties over a time scale longer than the transient detachment and re-attachment of each connector.

**Figure 4:**
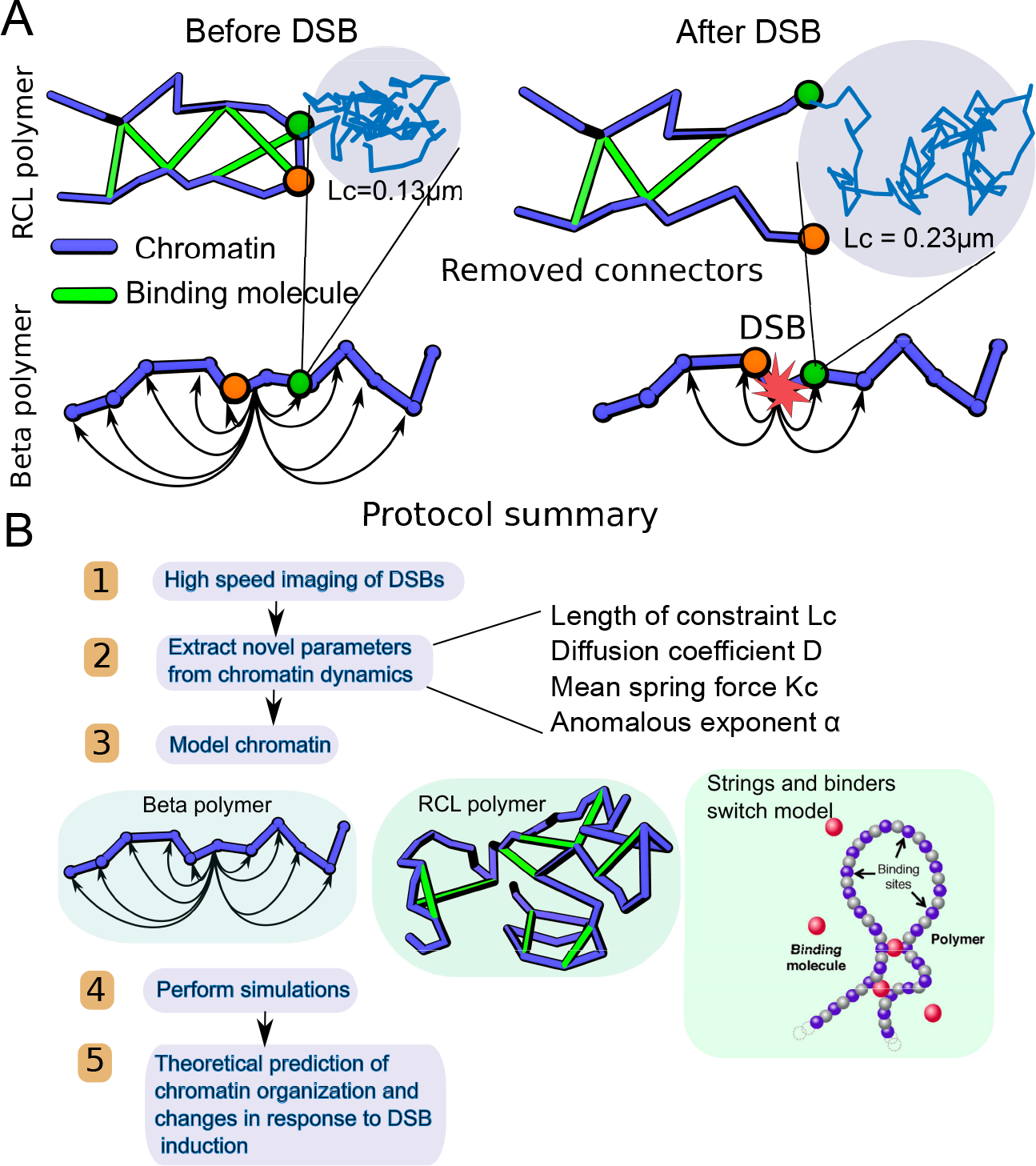
Chromatin modification represented by polymer models before and after DSB. **A.** An increase of *Lc* following DSB can be obtained by removing connectors (green) in RCL polymer model (top) or by removing long-range forces in the beta polymer model (bottom). **B.** Summary of the general procedure of converting information contained in the four parameters extracted from SPT of chromatin loci into a polymer representation. This polymer can be used to generate statistics not contained in the original trajectories.

### 4.2 Data calibration of cross-linkers

How CTCF and Cohesin or Condensin contribute to chromatin organization? This question can be explored using polymer model with a calibrated number of added connectors? A first calibration came from HiC data based on a polymer modeling [63]: this approach converts encounter probability of the HiC data into a short-range (intra-TAD organization) and long-range connectors. From this reconstruction approach, it is possible to compute a synthetic encounter probability map, which appears to be very similar to the empirical HiC one, suggesting that the reconstructed polymer model is an adequate average representation of chromatin organization at the scale of the HiC data. Once such a polymer is constructed, it is possible to explore transient properties such as the time and the probability for two loci to meet before a third one, and so one. Thus this analysis reveal the local chromatin exploration properties.

Another possible calibration is to determine the number of cross-linkers from local confined area, explored by a locus before and after the induction of DSB. In that case, when the length of confinement *L*_*c*_ increasing from 0.13 (before) to 0.23 *μm* (after) DSB [7], then the number of connectors needed for a RCL-polymer with a total of *N* = 100 monomers vary from *N*_*c*_ = 130 to 125, respectively (Fig. 4A).

Interestingly, if a large number of cross-linkers is required to condense a polymer to a small blob, surprisingly only ~ 4% are removed to match the decondensed DSB phase. In addition, the mean radius of gyration 15 〈*R*_*g*_〉 ≈150nm is mostly unchanged between the unbroken locus and DSB. However, the encounter time *τ*_*E*_ (defined above, see also Fig. 4D) changes from 1 to 2.8s, showing that removing few connectors could affect the local encounter time. Changing the number of connectors appears in the mean maximal distance between the two monomers, increasing from 0.33*μm* in the unbroken case to 0.75*μm* after DSB. After separation of the two strands, the remaining connectors maintain the two extremities with a maximal distance that decreased with *N*_*c*_ (Fig. 4A) left), showing that added random connectors serve to confine loci local environment, and prevent the two broken ends from drifting apart. This situation can facilitate re-ligation by non-homologous-end-joining (NHEJ), as well as locally opening the chromatin environment to allow access for the accumulation of repair proteins. To conclude, randomly positioned cross-linkers are used to model chromatin condensation and dynamics. Their minimum number can be adjusted to match experimental data of SPTs and HiC, offering a simple tool to represent chromatin organization at a given spatial scale, that can be used to explore the local chromatin environment and dynamical properties.

## 5 Available chromatin polymer packages

There are different types of available packages to compute and analyse the properties of loci mentioned in the past section: 1) Extract parameters from SPTs data 2) Reconstruct chromatin from HiC 3) Simulate polymer dynamics; single loci, compute MSD, 4) Reconstruct polymer model with specific properties such as *β*-polymer, accounting for viscosity or RCL.

### 5.1 Type 1

Matlab codes to compute the four parameters from SPTs [28, 7] are available at *http://bionewmetrics.org/analysis-of-spts/*. Fast algorithms of RCL-polymers [60] are available at *http://bionewmetrics.org/randomlycrosslinkedpolymer//*. Trajectories and polymer can be visualized in 3D and quantities such as mean-square-radius of gyration (MSRG), encounter probability (EP), and variance of the distance between loci, the Mean square displacement (MSD) and mean first encounter time (MFET) between loci are computed. Extension of this framework to include a single and multiple DSB, is provided in a Julia framework at http://bionewmetrics.org/randomly-cross-linked-polymer-following-dsb/.

### 5.2 Types 2-3

The numerical code to generate a generalized RCL polymer framework, consisting of two interacting Topologically Associating Domains (TADs) [63] is available at http://bionewmetrics.org/reconstruction-of-a-polymer-model-from-the-chromatin-capture-data/. Short random and fixed long-range connectors are added between TADs. Chromatin parameters can be calibrated directly from chromatin conformation capture data (3C, 5C, HiC) given TAD boundaries positions. Peaks within and between TADs lcoated in the encounter probability matrix are detected and converted in a number *N*_*c*_ of connectors. This framework enables to study the influence of short-range random connectors within TADs and long-range fixed connectors within and between TADs on the dynamic properties of loci in cross-linked chromatin.

### 5.3 Type 3-4

Fast numerical polymer simulations, incorporating internal and external forces, such as those from nuclear boundaries and diffusing molecules and structural proteins (CTCF) are implemented in an open source platform OpenMM [20] (*http://openmm.org/*). OpenMM framework is used in [19] to simulate polymers as a model of chromatin in confined nano-domains. Coarse-grained polymer simulations consisting of transient genomic loops with OpenMM are available at *https://mimakaev.bitbucket.io/index.html*.

A general molecular dynamics computational framework is LAMMPS [54] (https://lammps.sandia.gov). LAMMPS provides a flexible interface and packages to simulate polymer in confined domains. LAMMPS could also be used to simulate realizations of many types of molecular dynamics scenarios, such as blood flow in capillaries. LAMMPS was used in [72, 59, 14] to study the folding landscape of chromatin calibrated from CC contact data.

Reconstruction of a three-dimensional (3D) chromatin structures from ensemble CC contact maps remains a challenging task [55]. Models that suggest a solution to this problem take as an input CC contact maps and output an averaged (consensus) 3D structures by employing optimization algorithms to convert loci-loci encounter probability to a spatial distance, commonly based on the expected decay of the CC encounter probability with genomic distance [71]. In the model developed in [51], the average loci distances are computed from CC maps and additional constraints on chromatin folding from the expression of epigenomic markers see also [72].

3D structure reconstruction from a single cell HiC data was developed in [64] based on NMR principle, codes are available at (*https://github.com/TheLaueLab/nucdynami* Although average distances between genomic loci can be computed from the 3D structure, the dynamics of the reconstructed chromatin cannot be retrieved directly from the resulting structure because the final structure does not include information about cross-linkers, their distribution and dynamics. Bio-informatic tools [8] provide algorithms to extract parameters directly from CC contact maps, such as TAD boundaries and peak (see [56]), which enables to extract the position of fixed long-range connectors. The parameters extracted can be used for polymer simulations.

## 6 Conclusions

As the world of chromatin molecular biology and biophysics become more entangled it is essential to build a set of reliable tools to easily analyze biological data. Here we have described techniques and tools to study SPT data as well as how to use polymer models to interpret these results in the context of chromatin sub-organization. We further presented how to extract information from correlated loci analysis and how to interpret the changes in parameter values when the sampling rate Δ*t* is changing.

Models are a step towards a unified tool of SPT data with data generated by chromatin conformation capture techniques such as HiC (Fig. 4B). Such a tool would be immensely useful as it would take into account physical dynamics of chromatin at a local level while also giving an overview of interactions throughout globally. This type of unified model would allow us to better understand a number of complicated biological processes such as enhancer / promoter organization during transcriptional reprogramming, homology search during DSB repair and CRISPR/Cas9 editing. In particular, simulating the effects of integrating viral DNA on the chromatin environment would give new insight into how viruses cause mutations and ultimately disease.

## References

[1] Harsha Agarwal, Matthias Reisser, Celina Wortmann, and J Christof M Gebhardt. Direct observation of cell-cycle-dependent interactions between ctcf and chromatin. Biophysical journal, 112(10):2051–2055, 2017.

[2] A. Amitai, I. Kupka, and D. Holcman. Computation of the mean first-encounter time between the ends of a polymer chain. Phys. Rev. Lett, 109:108302, 2012.

[3] A Amitai, M Toulouze, K Dubrana, and D Holcman. Analysis of single locus trajectories for extracting in vivo chromatin tethering interactions. PLoS Computational Biology, 11(8):e1004433, 2015.

[4] A. Amitai, M. Toulouze, K. Dubrana, and D. Holcman. Extracting in vivo interactions acting on the chromatin from a statistical analysis of single locus trajectories. Personal Communication, 2015.

[5] Assaf Amitai and David Holcman. Polymer model with long-range interactions: Analysis and applications to the chromatin structure. Physical Review E, 88(5):052604, 2013.

[6] Assaf Amitai and David Holcman. Polymer physics of nuclear organization and function. Physics Reports, 678:1–83, 2017.

[7] Assaf Amitai, Andrew Seeber, Susan M Gasser, and David Holcman. Visualization of chromatin decompaction and break site extrusion as predicted by statistical polymer modeling of single-locus trajectories. Cell Reports, 18(5):1200–1214, 2017.

[8] Ferhat Ay and William S Noble. Analysis methods for studying the 3d architecture of the genome. Genome biology, 16(1):183, 2015.

[9] Mariano Barbieri, Mita Chotalia, James Fraser, Liron-Mark Lavitas, Josée Dostie, Ana Pombo, and Mario Nicodemi. Complexity of chromatin folding is captured by the strings and binders switch model. Proceedings of the National Academy of Sciences, 109(40):16173–16178, 2012.

[10] M. Bohn, D. W. Heermann, and R. van Driel. Random loop model for long polymers. Phys. Rev. E, 76:051805, 2007.

[11] Stanislav Burov, S. M. Ali Tabei, Toan Huynh, Michael P. Murrell, Louis H. Philipson, Stuart A. Rice, Margaret L. Gardel, Norbert F. Scherer, and Aaron R. Dinner. Distribution of directional change as a signature of complex dynamics. Proceedings of the National Academy of Sciences, 110(49):19689–19694, 2013.

[12] Baohui Chen, Juan Guan, and Bo Huang. Imaging specific genomic dna in living cells. Annual review of biophysics, 45:1–23, 2016.

[13] Hongtao Chen, Michal Levo, Lev Barinov, Miki Fujioka, James B Jaynes, and Thomas Gregor. Dynamic interplay between enhancer– promoter topology and gene activity. Nature genetics, 50(9):1296, 2018.

[14] Michele Di Pierro, Ryan R Cheng, Erez Lieberman Aiden, Peter G Wolynes, and José N Onuchic. De novo prediction of human chromosome structures: Epigenetic marking patterns encode genome architecture. Proceedings of the National Academy of Sciences, page 201714980, 2017.

[15] David Dickerson, Marek Gierliński, Vijender Singh, Etsushi Kitamura, Graeme Ball, Tomoyuki U Tanaka, and Tom Owen-Hughes. High resolution imaging reveals heterogeneity in chromatin states between cells that is not inherited through cell division. BMC Cell Biology, 17(1):33, 2016.

[16] V. Dion and S. M. Gasser. Chromatin movement in the maintenance of genome stability. Cell, 152:1355–1364, 2013.

[17] Vincent Dion, Véronique Kalck, Andrew Seeber, Thomas Schleker, and Susan M Gasser. Cohesin and the nucleolus constrain the mobility of spontaneous repair foci. EMBO Reports, 14(11):984, 2013.

[18] M Doi and SF Edwards. The Theory of Polymer Dynamics Clarendon. Oxford, 1986.

[19] Boryana Doyle, Geoffrey Fudenberg, Maxim Imakaev, and Leonid A Mirny. Chromatin loops as allosteric modulators of enhancer-promoter interactions. PLoS computational biology, 10(10):e1003867, 2014.

[20] Peter Eastman, Jason Swails, John D Chodera, Robert T McGibbon, Yutong Zhao, Kyle A Beauchamp, Lee-Ping Wang, Andrew C Simmonett, Matthew P Harrigan, Chaya D Stern, et al. Openmm 7: Rapid development of high performance algorithms for molecular dynamics. PLoS computational biology, 13(7):e1005659, 2017.

[21] Brian P English, Vasili Hauryliuk, Arash Sanamrad, Stoyan Tankov, Nynke H Dekker, and Johan Elf. Single-molecule investigations of the stringent response machinery in living bacterial cells. Proceedings of the National Academy of Sciences, 108(31):E365–E373, 2011.

[22] Nicholas A Frost, Hari Shroff, Huihui Kong, Eric Betzig, and Thomas A Blanpied. Single-molecule discrimination of discrete perisynaptic and distributed sites of actin filament assembly within dendritic spines. Neuron, 67(1):86–99, 2010.

[23] J Christof M Gebhardt, David M Suter, Rahul Roy, Ziqing W Zhao, Alec R Chapman, Srinjan Basu, Tom Maniatis, and X Sunney Xie. Single-molecule imaging of transcription factor binding to dna in live mammalian cells. Nature methods, 10(5):421, 2013.

[24] Rodolfo Ghirlando and Gary Felsenfeld. Ctcf: making the right connections. Genes & development, 30(8):881–891, 2016.

[25] Emmanuelle Guillou, Arkaitz Ibarra, Vincent Coulon, Juan Casado-Vela, Daniel Rico, Ignacio Casal, Etienne Schwob, Ana Losada, and Juan Méndez. Cohesin organizes chromatin loops at dna replication factories. Genes & development, 24(24):2812–2822, 2010.

[26] Judith HI Haarhuis, Robin H van der Weide, Vincent A Blomen, J Omar Yéñez-Cuna, Mario Amendola, Marjon S van Ruiten, Peter HL Krijger, Hans Teunissen, René H Medema, Bas van Steensel, et al. The cohesin release factor wapl restricts chromatin loop extension. Cell, 169(4):693–707, 2017.

[27] Anders S Hansen, Iryna Pustova, Claudia Cattoglio, Robert Tjian, and Xavier Darzacq. Ctcf and cohesin regulate chromatin loop stability with distinct dynamics. Elife, 6:e25776, 2017.

[28] Michael H Hauer, Andrew Seeber, Vijender Singh, Raphael Thierry, Ragna Sack, Assaf Amitai, Mariya Kryzhanovska, Jan Eglinger, David Holcman, Tom Owen-Hughes, et al. Histone degradation in response to dna damage enhances chromatin dynamics and recombination rates. Nature Structural & Molecular Biology, 2017.

[29] Dieter W Heermann. Physical nuclear organization: loops and entropy. Current Opinion in Cell Biology, 23(3):332–337, 2011.

[30] Patrick Heun, Thierry Laroche, Kenji Shimada, Patrick Furrer, and Susan M Gasser. Chromosome dynamics in the yeast interphase nucleus. Science, 294(5549):2181–2186, 2001.

[31] D. Holcman, N. Hoze, and Z. Schuss. Narrow escape through a funnel and effective diffusion on a crowded membrane. Phys Rev E, 84:021906, 2011.

[32] D Holcman, Nathanaël Hoze, and Z Schuss. Analysis and interpretation of superresolution single-particle trajectories. Biophysical journal, 109(9):1761–1771, 2015.

[33] David Holcman, Pierre Parutto, Joseph E Chambers, Marcus Fantham, Laurence J Young, Stefan J Marciniak, Clemens F Kaminski, David Ron, and Edward Avezov. Single particle trajectories reveal active endoplasmic reticulum luminal flow. Nature cell biology, 20(10):1118, 2018.

[34] David Holcman, Pierre Parutto, Joseph E Chambers, Marcus Fantham, Laurence J Young, Stefan J Marciniak, Clemens F Kaminski, David Ron, and Edward Avezov. Single particle trajectories reveal active endoplasmic reticulum luminal flow. Nature cell biology, 20(10):1118, 2018.

[35] N. Hozé and D. Holcman. Coagulation-fragmentation for a finite number of particles and application to telomere clustering in the yeast nucleus. Phys. Lett. A, 845:5043, 2012.

[36] Nathanael Hoze and David Holcman. Residence times of receptors in dendritic spines analyzed by stochastic simulations in empirical domains. Biophysical journal, 107(12):3008–3017, 2014.

[37] Nathanaël Hoze and David Holcman. Recovering a stochastic process from super-resolution noisy ensembles of single-particle trajectories. Physical Review E, 92(5):052109, 2015.

[38] Nathanaël Hozé and David Holcman. Statistical methods for large ensembles of super-resolution stochastic single particle trajectories in cell biology. Annual Review of Statistics and Its Application, 4:189–223, 2017.

[39] Nathanaël Hozé and David Holcman. Statistical methods for large ensembles of super-resolution stochastic single particle trajectories in cell biology. Annual Review of Statistics and Its Application, 4:189–223, 2017.

[40] Ignacio Izeddin, Vincent Récamier, Lana Bosanac, Ibrahim I Cissé, Lydia Boudarene, Claire Dugast-Darzacq, Florence Proux, Olivier Bénichou, Raphaël Voituriez, Olivier Bensaude, et al. Single-molecule tracking in live cells reveals distinct target-search strategies of transcription factors in the nucleus. Elife, 3:e02230, 2014.

[41] DA Jackson, P Dickinson, and PR Cook. The size of chromatin loops in hela cells. The EMBO Journal, 9(2):567–571, 1990.

[42] Eldad Kepten, Irena Bronshtein, and Yuval Garini. Improved estimation of anomalous diffusion exponents in single-particle tracking experiments. Physical Review E, 87(5):052713, 2013.

[43] Eldad Kepten, Aleksander Weron, Grzegorz Sikora, Krzysztof Burnecki, and Yuval Garini. Guidelines for the fitting of anomalous diffusion mean square displacement graphs from single particle tracking experiments. PLoS One, 10(2):e0117722, 2015.

[44] Tae Hoon Kim, Ziedulla K Abdullaev, Andrew D Smith, Keith A Ching, Dmitri I Loukinov, Roland D Green, Michael Q Zhang, Victor V Lobanenkov, and Bing Ren. Analysis of the vertebrate insulator protein ctcf-binding sites in the human genome. Cell, 128(6):1231–1245, 2007.

[45] Thomas J Lampo, Andrew S Kennard, and Andrew J Spakowitz. Physical modeling of dynamic coupling between chromosomal loci. Biophysical journal, 110(2):338–347, 2016.

[46] Zhe Liu, Luke D Lavis, and Eric Betzig. Imaging live-cell dynamics and structure at the single-molecule level. Molecular cell, 58(4):644–659, 2015.

[47] Philipp G Maass, A Rasim Barutcu, David M Shechner, Catherine L Weiner, Marta Melé, and John L Rinn. Spatiotemporal allele organization by allele-specific crispr live-cell imaging (snp-cling). Technical report, Nature Publishing Group, 2018.

[48] Suliana Manley, Jennifer M Gillette, George H Patterson, Hari Shroff, Harald F Hess, Eric Betzig, and Jennifer Lippincott-Schwartz. High-density mapping of single-molecule trajectories with photoactivated localization microscopy. Nature methods, 5(2):155, 2008.

[49] Ralf Metzler and Joseph Klafter. The restaurant at the end of the random walk: recent developments in the description of anomalous transport by fractional dynamics. Journal of physics. A, mathematical and general, 37(31):R161–R208, 2004.

[50] Frank R Neumann, Vincent Dion, Lutz R Gehlen, Monika Tsai-Pflugfelder, Roger Schmid, Angela Taddei, and Susan M Gasser. Targeted ino80 enhances subnuclear chromatin movement and ectopic homologous recombination. Genes & development, 26(4):369–383, 2012.

[51] Juan D Olarte-Plata, Noelle Haddad, Cédric Vaillant, and Daniel Jost. The folding landscape of the epigenome. Physical biology, 13(2):026001, 2016.

[52] Roxanne Oshidari, Jonathan Strecker, Daniel KC Chung, Karan J Abraham, Janet NY Chan, Christopher J Damaren, and Karim Mekhail. Nuclear microtubule filaments mediate non-linear directional motion of chromatin and promote dna repair. Nature communications, 9, 2018.

[53] R. W. Pastor, R. Zwanzig, and A. Szabo. Diffusion limited first contact of the ends of a polymer: Comparison of theory with simulation. J. Chem. Phys., 105:3878–3882, 1996.

[54] Steve Plimpton. Fast parallel algorithms for short-range molecular dynamics. Journal of computational physics, 117(1):1–19, 1995.

[55] Angelo R. and Christophe Z. Chapter nine - computational models of large-scale genome architecture. In New Models of the Cell Nucleus: Crowding, Entropic Forces, Phase Separation, and Fractals, volume 307 of International Review of Cell and Molecular Biology, pages 275 – 349. Academic Press, 2014.

[56] Suhas SP Rao, Miriam H Huntley, Neva C Durand, Elena K Stamenova, Ivan D Bochkov, James T Robinson, Adrian L Sanborn, Ido Machol, Arina D Omer, Eric S Lander, et al. A 3d map of the human genome at kilobase resolution reveals principles of chromatin looping. Cell, 159(7):1665–1680, 2014.

[57] Andrew Seeber, Michael H Hauer, and Susan M Gasser. Chromosome dynamics in response to dna damage. Annual review of genetics, (0), 2018.

[58] Andrew Seeber, Michael H Hauer, and Susan M Gasser. Chromosome dynamics in response to dna damage. Annual review of genetics, (0), 2018.

[59] Guang Shi, Lei Liu, Changbong Hyeon, and Dave Thirumalai. Interphase human chromosome exhibits out of equilibrium glassy dynamics. Nature communications, 9(1):3161, 2018.

[60] O Shukron and D Holcman. Statistics of randomly cross-linked polymer models to interpret chromatin conformation capture data. Physical Review E, 96(1):012503, 2017.

[61] Ofir Shukron, Michael Hauer, and David Holcman. Two loci single particle trajectories analysis: constructing a first passage time statistics of local chromatin exploration. Scientific reports, 7(1):10346, 2017.

[62] Ofir Shukron and David Holcman. Heterogeneous cross-linked polymers to reconstruct chromatin reorganization during cell differentiation. bioRxiv, page 235051, 2017.

[63] Ofir Shukron and David Holcman. Transient chromatin properties revealed by polymer models and stochastic simulations constructed from chromosomal capture data. PLOS Computational Biology, 13(4):e1005469, 2017.

[64] Tim J Stevens, David Lando, Srinjan Basu, Liam P Atkinson, Yang Cao, Steven F Lee, Martin Leeb, Kai J Wohlfahrt, Wayne Boucher, Aoife OShaughnessy-Kirwan, et al. 3d structures of individual mammalian genomes studied by single-cell hi-c. Nature, 544(7648):59, 2017.

[65] Johannes Stigler, Gamze Ö Çamdere, Douglas E Koshland, and Eric C Greene. Single-molecule imaging reveals a collapsed conformational state for dna-bound cohesin. Cell reports, 15(5):988–998, 2016.

[66] Terence Strick, Jean-François Allemand, Vincent Croquette, and David Bensimon. Twisting and stretching single dna molecules. Progress in biophysics and molecular biology, 74(1-2):115–140, 2000.

[67] TR Strick, MN Dessinges, G Charvin, NH Dekker, JF Allemand, D Bensimon, and V Croquette. Stretching of macromolecules and proteins. Reports on Progress in Physics, 66(1):1, 2002.

[68] A. Taddei and S. M. Gasser. Structure and function in the budding yeast nucleus. Genetics, 192:107–29, 2012.

[69] Guido Tiana, Assaf Amitai, Tim Pollex, Tristan Piolot, David Holcman, Edith Heard, and Luca Giorgetti. Structural fluctuations of the chromatin fiber within topologically associating domains. Biophysical journal, 110(6):1234–1245, 2016.

[70] Jolien Suzanne Verdaasdonk, Paula Andrea Vasquez, Raymond Mario Barry, Timothy Barry, Scott Goodwin, M Gregory Forest, and Kerry Bloom. Centromere tethering confines chromosome domains. Molecular cell, 52(6):819–831, 2013.

[71] Siyu Wang, Jinbo Xu, and Jianyang Zeng. Inferential modeling of 3d chromatin structure. Nucleic acids research, 43(8):e54–e54, 2015.

[72] Bin Zhang and Peter G Wolynes. Topology, structures, and energy landscapes of human chromosomes. Proceedings of the National Academy of Sciences, 112(19):6062–6067, 2015.

[73] Alexandra Zidovska, David A Weitz, and Timothy J Mitchison. Micronscale coherence in interphase chromatin dynamics. Proceedings of the National Academy of Sciences, page 201220313, 2013.

